# Assessing Structural Prediction Accuracy for Nanobody–Small Molecule Complexes

**DOI:** 10.64898/2025.12.12.693880

**Authors:** Berta Bori-Bru, Juan Pablo Salvador, Ramon Crehuet

## Abstract

Generative models have advanced profusely in recent years, generating great impact in computational modelling and structural bioinformatics. A part of computational protein design focuses on antibody and nanobody engineering, targeting proteic epitopes. By contrast, the development of nanobodies for small-molecule sensing remains a largely experimental field. In this work, we evaluate the performance of several state-of-the-art prediction softwares (AlfaFold3, Chai-1, Boltz-1, RosettaFold-AllAtom, FlowDock and OmegaFold), on nanobody-small molecule complexes, in an effort to pave the way of computational studies in this area. We tested the general nanobody and CDR accuracy, as well as ligand placement and orientation. We also explored the correlation of the results with intrinsic metrics of the models, such as pLDDT, and with the number of samples and recycles. Results show that most predictors perform well at predicting nanobody structures, but some struggle at ligand placement. AlphaFold3 outperformed the other softwares in all these tasks. Co-folding increased the accuracy of the predictions, modelling better CDR1. Contact analysis revealed that CDR1 was mostly involved in ligand binding instead of CDR3. This could point to a memorization tendency in the models, as most nanobody-antigen complexes target other proteins. Results also pointed to pLDDT as a good score to indicate CDR accuracy. Accuracy did slightly improve when increasing the number of samples but did not with the number of recycles. These findings highlight both the advantages and limitations of structure prediction methods for nanobody–small molecule complexes.

## Introduction

Protein prediction models have evolved at a high speed in recent years. From the first monomer predictions of AlphaFold in 2021^1^ to the recent development of AlphaFold3^2^ and other diffusion-based deep learning softwares^3,4^, only three years have elapsed. Now, predicting structures containing combinations of proteins, nucleic acids, and small molecules is possible. Structure prediction algorithms have shaped the way we study structural biology, which led to the bestowal of the 2024 Chemistry Nobel Prize to their main contributors^5^. The structure prediction revolution triggered a paradigm shift in protein design^6,7^. Once experimentally costly, this task is now greatly streamlined by computational methods, though experimental validation remains indispensable.

The design of protein binders for small-molecules has flourished as a result of the protein design revolution. It has led to the creation of biosensors^8,9^, therapeutic compounds^10,11^ and even enzymes^12–14^. Most of these new protein structures are based on artificial folds developed by protein design groups. However, to design highly selective and affine binders, nature relies on antibodies and their smaller “cousins”, nanobodies.

Nanobodies (Nbs) were discovered in 1993 in the laboratory of Richard Hammers, where they observed camelid antibodies containing only a heavy chain, and not the light and heavy chains found in conventional antibodies^15^. Conventional antibodies are composed of four polypeptidic chains, two light and two heavy chains, which have a total molecular weight of 150 kDa. Nanobodies, or also so-called single-chain antibodies, are the recombinant part of the variable fraction of the heavy chain antibodies. The fact that Nbs are composed by only one polypeptidic chain confers an additional stability, maintaining the affinity constant against their corresponding antigen. According to their low molecular weight (12-15 kDa), Nbs can be internalized in cells^16^, and, due to their recombinant nature, can be engineered in different fusion proteins^17^.

In comparison to conventional antibodies, they only contain half of the complementary determining regions (CDRs) that are responsible for the specificity and selectivity against the target analyte. CDRs are disordered regions composed by 7-17 amino acids that form the high affinity binding region and source of variability of both antibodies and nanobodies. The other more rigid and conserved regions are named framework regions (FWRs). Despite their small size, nanobodies are on par with antibodies in selectivity and specificity, reaching nanomolar or even picomolar ranges of affinity^18^. Although the vast majority of nanobodies are used to target proteins, they have also been employed to create immunosensors to target small molecules, thanks to their outstanding properties^19–22^. Despite the computational advances in the last years, most of the efforts in the field have been experimental and the few computational advances were more focused on sequence-based approaches^23–25^, with only one structural work published^26^ in 2025.

Meanwhile, there have been attempts in the field of computational structural biology to benchmark structure predictors for protein-ligand compounds^27–29^, albeit not focused on nanobody or antibody usage; and nanobody and antibody benchmarks have overlooked small-molecule binding^30,31^. This might be due to the low availability of structures to produce a benchmarking: in SabDab database^32^ only 3% of antibodies and 1% of nanobodies are bound to small-molecules. However, the potential gains from this could result in significant advancements in the field.

In this study we present a benchmarking on the performance of different state-of-the-art structure prediction softwares on different nanobody-small molecule analyte systems. We analyse their accuracy on protein prediction and ligand placement, as well as CDR prediction and their relationship with binding contacts. We also assess how this accuracy is related to the reported confidence scores for the best couple of methods and the dependence of the results in the number of recycles and diffusion samples.

## Results

The softwares used in this benchmark include folding programs like AlphaFold3 (AF3)^2^, Chai-1^3^, Boltz-1 and Boltz-1x^4^, as well as RosettaFold All-Atom (RFAA)^33^ and OmegaFold^34^. We also added the docking software FlowDock^35^ (see Methods).

In the benchmark, we decided to test the capabilities of all the aforementioned softwares for the prediction of the nanobody-only as well as the co-folded complex (nanobody and small-molecule) structures for ten complexes retrieved from SAbDab^32^ (see Methods). We also compare how the quality of the predictions changes when the predicted sequences are in the training sets of the predictors.

### Co-folding increases nanobody structure accuracy

To assess the overall accuracy of the predicted nanobody structures we calculated the RMSD of the Ca between the crystal structures and the predictions, both with and without co-folding. In Figure 1A we observe that all predictors achieve satisfactory results, with RMSDs below 1.7Å. The best predictors for this dataset are AF3 and Chai-1 when co-folding the nanobody and the small molecule, providing similar results between them, but significantly better than the rest of predictors. Both Boltz-1 and Boltz-1x generate similar structures. This is expected as Boltz-1x was mainly developed to avoid unphysical poses of the ligand and there seems to be no clashes already in the predicted structures by Boltz-1. FlowDock provides the worst results among the compared softwares. Even when prompted with a structure from Chai-1 as template, it is not able to reproduce the same accuracy as Chai-1. This could be due to the focus of FlowDock on blind docking and not structure prediction. RFAA provides decent results when only folding the protein. OmegaFold produces similar results as RFAA, at a higher speed. However, the precision obtained using co-folding surpasses the benefits of lower computational cost, rendering OmegaFold less recommendable for this task.

**Figure 1:**
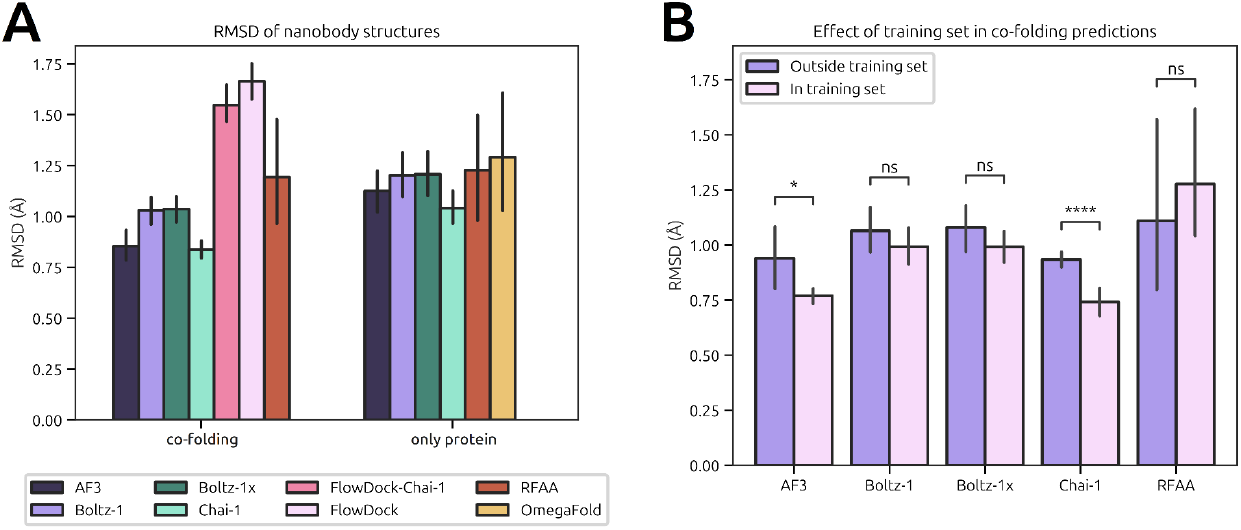
Comparison of protein structure accuracy for different predictors. **A**: Comparison between nanobody RMSD in co-folding predictions versus only protein ones. **B**: Effect of the presence of the complexes in the training set of the predictors to the RMSD of co-folded structures.

Ligand co-folding increases the predicted structure quality for the majority of methods. Such increment is stronger for Chai-1 and AF3, and does not appear in RFAA, meaning its predictions are not affected by co-folding.

As the benchmarking dataset contains some structures that were used in the training sets for these deep learning predictors, we analysed whether the predicted structures were more accurate if seen during the training phase. Figure S2 shows that the structure predictions improve when the queried sequence was used for training, but the differences are minor and not significant. Surprisingly, FlowDock performance worsens when the sequence was included in the training set.

Including the effect of co-folding to this analysis, in Figure 1B only AF3 and Chai-1 provide significantly better results when predicting structures in their training set. Thus, we can expect slightly worse results when querying unknown sequences, as is done in protein design.

### Accurate binding site location, but poor ligand orientation

Having studied the accuracy of the protein structure, the next analysis assesses the accuracy of ligand positioning. For that, we calculated the center of mass distance between the predicted and crystal ligands, and the RMSD of ligand atoms to account for ligand orientation. Usually, the process of docking the ligand in a protein structure is uncoupled from the structure prediction problem, but due to the CDRs flexibility, we expected the presence of the ligand to stabilize conformations that are missing or unstable in the apo structure.

In Figure 2A, we observe that the predictors place the ligands with a deviation of 5Å from the center of mass, which indicates a moderate accuracy in pose predictions. For the ligand RMSD, the overall results show values of 8Å, reflecting poor ligand orientation capabilities. AF3 provides the most accurate results, outperforming the other methods in these tasks. It predicts binding site distances of 2Å, indicating a much better ligand placement; and RMSD of 6Å, which is still moderate in orientation accuracy. Chai-1 is the second best program, and RFAA provides similar results to the other predictors, but it tends to generate steric clashes between protein and ligand chains (see Figure S3) that render their predictions unreliable. FlowDock performs similarly to other predictors in ligand placement. Although a better performance might be expected since it is a dedicated docking program, its accuracy might be limited due to the poor accuracy in the predicted protein backbone.

**Figure 2:**
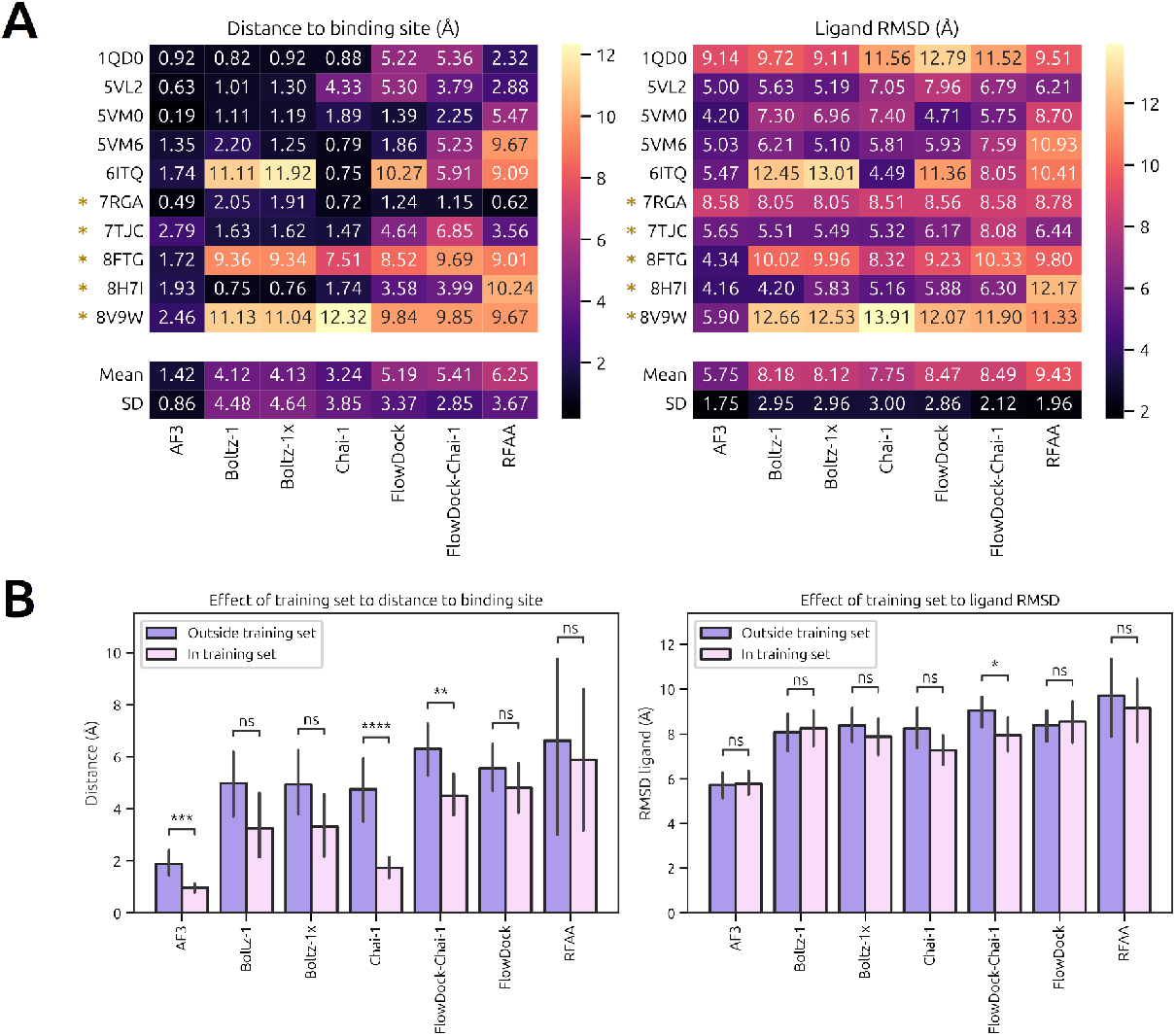
Ligand placement precision assessed by binding site distance (left column) and ligand RMSD (right column) for different predictors. **A**: Heatmap analysing the ligand placement for each system and each predictor. * indicate that the system is not in the training set of the predictors. **B**: Effect of the presence of the complexes in the training set of the predictors on ligand placement.

Analysing the individual nanobody cases in Figure 2A, we observe some interesting behaviours. 8FTG, 8V9W and 6ITQ trick all of the predictors except AF3 into placing the ligand in the wrong position. In 8FTG, the ligand is bound by two tyrosine residues through pi-stacking, placed next to CDR3. Only AF3 places the ligand in the correct spot, the other softwares position it outside of, but still near, CDR3. Similarly, for 8V9W all predictors except for AF3 place the ligand in CDR3 while its correct position is below CDR1. In 6ITQ, the ligand should be placed near CDR1 and is placed next to CDR3 instead by all predictors except for AF3 and Chai-1. We speculate that the cause of this phenomenon could be that most of the nanobodies bind to other proteins through CDR3. As these constitute most of the available nanobody structures, predictors might be predisposed to placing the ligands in CDR3. These results are coherent with the ones reported by Škrinjar and coworkers^28^, where they find that modern structure predictors are biased to previously known ligand poses.

Comparing binding site differences with RMSD, we also observe a couple of discrepancies in nanobodies 7RGA and 1QD0. For the former, this is due to a carboxylic tail from the methotrexate ligand, which is predicted in an elongated conformation and is folded unto itself in the crystal. For 1QD0, there are some differences in the alignment of the ligand, RR6, but that is to be expected as it is the biggest ligand in this benchmark, and thus has more rotable bonds, leading to high RMSD. However, the pose is mostly correct and the center of mass distance is rather low, indicating that it is placed in the correct binding site.

Results in Figure 2B hint to a relationship between the inclusion of a nanobody in the training set and the accuracy of the ligand placement. Regarding binding site distance, it seems that the complexes in the training sets have better ligand placements, but it is only significant for AF3 and Chai-1. This could be due to the low size of the benchmarking dataset. In contrast, when considering the RMSD, the predictions seem to be as accurate in nanobodies in the training set as for those outside of it. The high ligand RMSD highlights the challenge of predicting the ligand conformation when bound to the nanobody, especially for flexible ligands.

### CDR length determines accuracy

The previous results, specifically the Cα RMSD, provide an insight into the accuracy of the predictors for the protein structure. However, as seen in the last section, the binding process occurs mainly near the CDRs. Moreover, the framework regions (FWRs) follow a rigid β-sheet pattern and high sequence conservation. Therefore, one might expect to see less accuracy in the CDR RMSD and also to see more variation when including the ligand in the predictions. In Figure 3 we show the results for that analysis.

**Figure 3:**
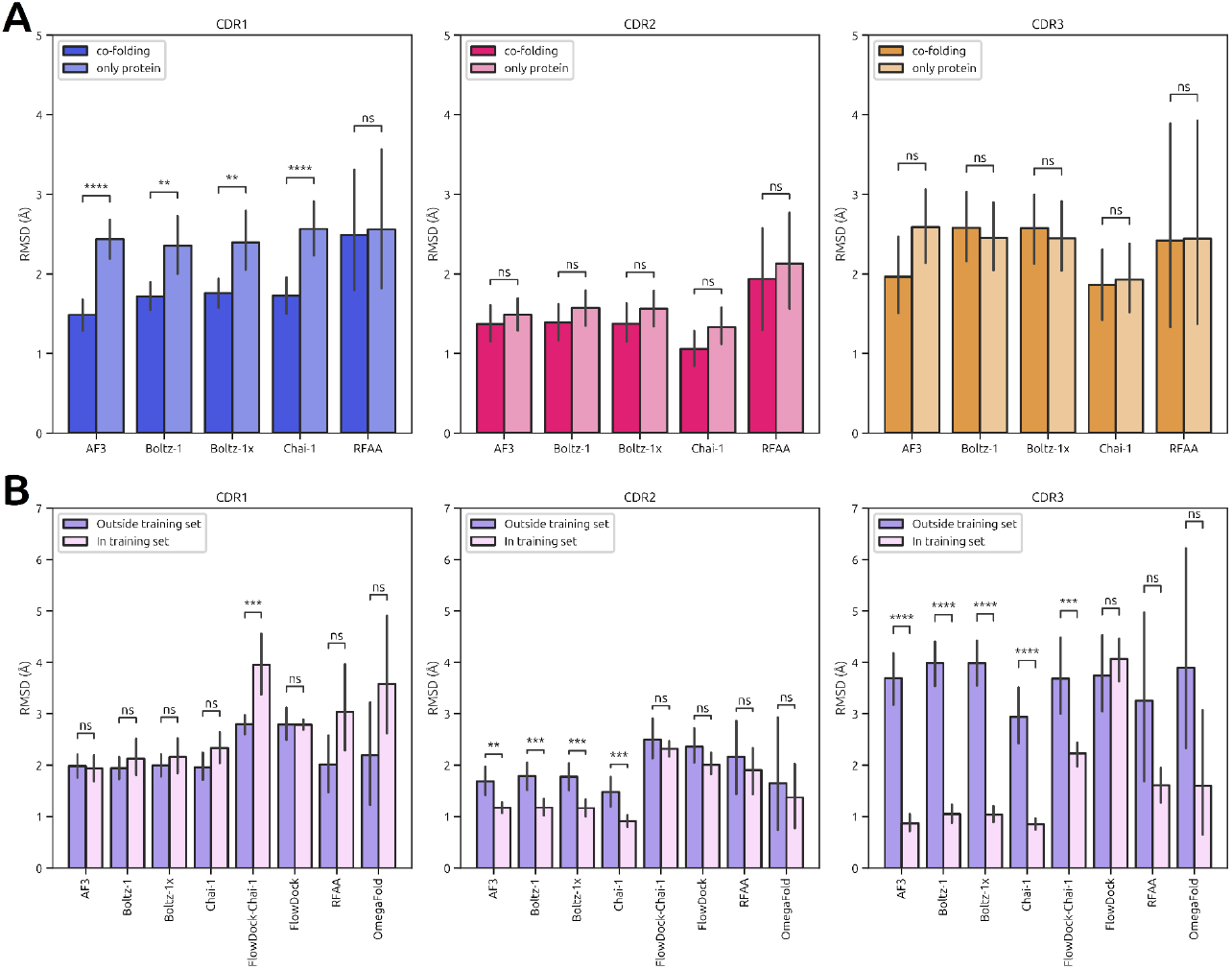
CDR RMSD for different structure predictors. **A**: Effect of co-folding on CDR predictions. **B**: Effect of inclusion in the training set on CDR predictions.

In Figure 3A, CDR3 has the highest RMSD score out of the 3 CDRs. This is to be expected since it commonly has the largest sequence, and thus is more flexible and difficult to model. CDR2 has the lowest RMSD, as it is also the shortest CDR.

Ligand co-folding significantly decreases the RMSD of CDR1, but not of CDR2 nor CDR3. In contrast, in Figure 3B the inclusion in the training set does not affect CDR1. This suggests that CDR1 plays an important role in binding small molecules and is more affected by ligand presence, barely noticing the effect of the inclusion in the training set. In contrast, accuracy in CDR2 and CDR3 is much greater for structures present in the training set of the predictors. This could be due to the intrinsic flexibility of these loop regions, being more difficult to model

### CDR1 is the main contributor to binding

To assess how much each CDR contributed to ligand binding, we calculated the number of contacts that each CDR and CDR neighbourhood (residues at 8Å from each atom in the CDR) had with the ligand (see Methods). As AF3 and Chai-1 are the best couple of predictors, from this section onwards, we will only report the results of these two methods.

The results in Figure 4A indicate that CDR1 has significantly more interactions with the ligand than CDR2 and CDR3, confirming our prior hypothesis. This is consistent for both AF3, Chai-1 and the crystal structures, with no significant differences between these three classes (Figure S5). This indicates that both of these structure predictors correctly capture the chemical nuances present in nanobody-small molecule systems.

**Figure 4:**
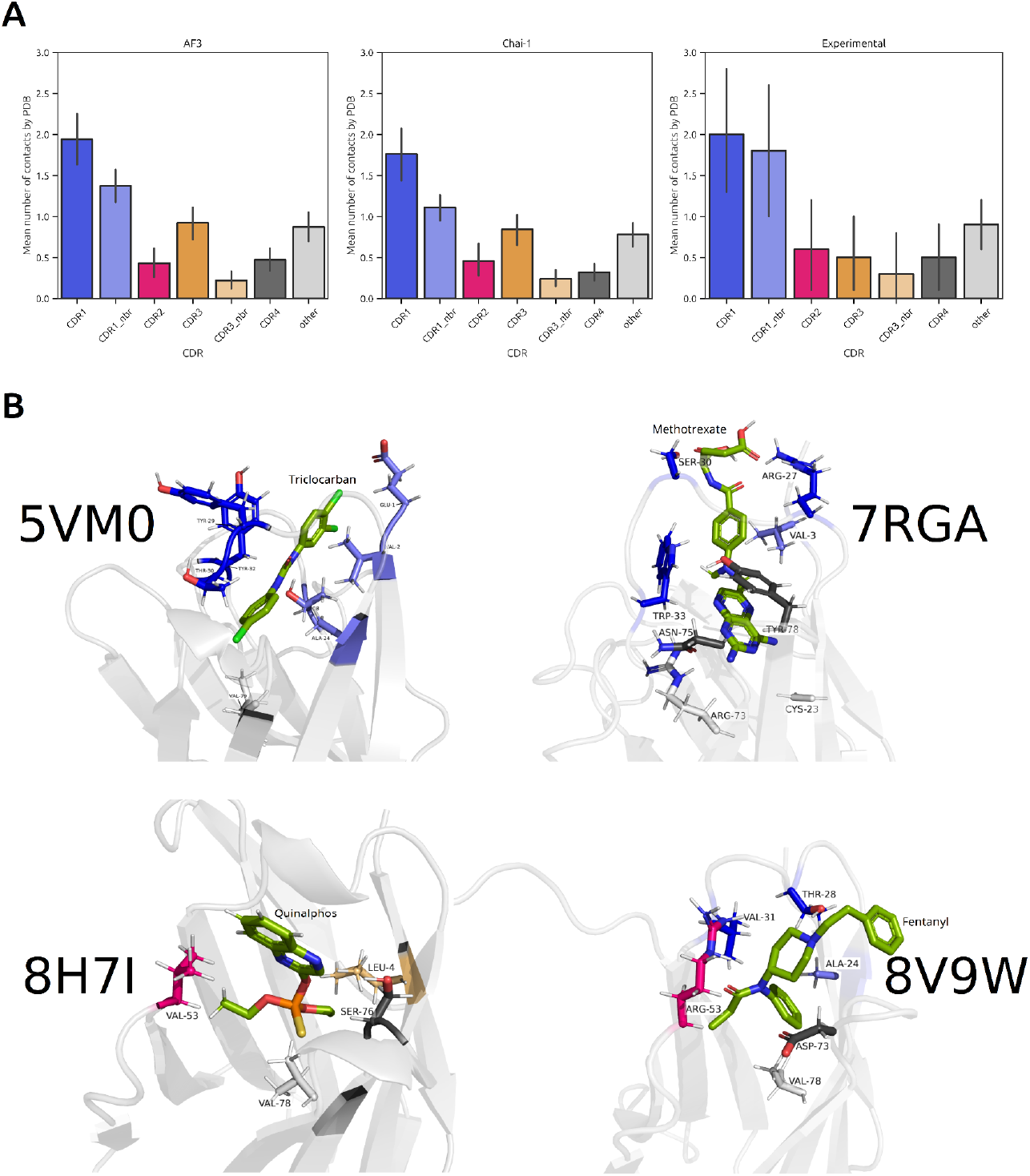
Analysis of the number of CDR and CDR neighbourhood (nbr) contacts. **A**: Number of contacts in predictions from AF3, Chai-1 and crystal PDB structures. **B**: Representations of the contacts of four nanobodies. Ligands in green, CDR1 contact residues in blue, CDR2 in magenta, CDR3 in yellow and CDR4 region in grey.

The CDR4 region plays a role as important as CDR2 in binding small molecules. It is a loop situated in FWR3, between CDR2 and CDR3^36,37^. Its importance was already noticed by Fanning and Horn^37^ for the nanobody 7RGA in its key role to binding methotrexate and even aiding in the stabilization of CDR1. Figure S6 indicates that 6ITQ, 8V9W, and 8H7I also have important contributions from the CDR4 region in binding cortisol, fentanyl and quinalphos, respectively.

In addition to the amino acids surrounding the CDRs, there are more faraway residues that also interact with the ligand (“other” column in Figure 4A). To check to which amino acids this field refers to, some structures with high contributions of “other” residues have been visually inspected using PyMOL (Figure 4B). Surprisingly, most of the “other” residues correspond to VAL78, just below the CDR4 region. This residue forms the bed of the binding site and has been described as a binding residue in quinalphos and fentanyl targeting nanobodies^38,39^. It might also be the case for ARG73 in 7RGA, which is not mentioned in the original publication. Furthermore, in some of the structures, we have found terminal residues that contribute to ligand binding, such as residue VAL4 in 7RGA (light blue) and triclocarban-targeting nanobodies (5VL2, 5VM6, 5VM0) or LEU4 in 8H7I (light yellow). These interactions were also described in the articles describing triclocarban and quinalphos (8H7I) targeting nanobodies^36,38^. This reinforces the idea that, in addition to the CDRs, other residues surrounding the ligand also play important roles in nanobody–epitope binding.

### Confidence scores correlate with structure accuracy

We next wondered if the predictor confidence parameters could inform about the quality of the model. These parameters, often measured with pLDDT, PTM and iPTM are of great interest when making blind predictions, as protein designs can be scored by them, if they are reliable.

Here we report the correlation of the pLDDT with the RMSD for each CDR only for AF3, as Chai-1 does not report the pLDDT of its predictions. Both for co-folding and protein-only cases we obtain a good correlation between pLDDT and CDR RMSD (Figure 5A). In accordance with previous results, co-folding not only increases the accuracy of CDR1, but it also increases its pLDDT score, as reported in previous articles^2,40^. This showcases the capacity of AF3 to self-assess the accuracy of the predicted structures.

**Figure 5:**
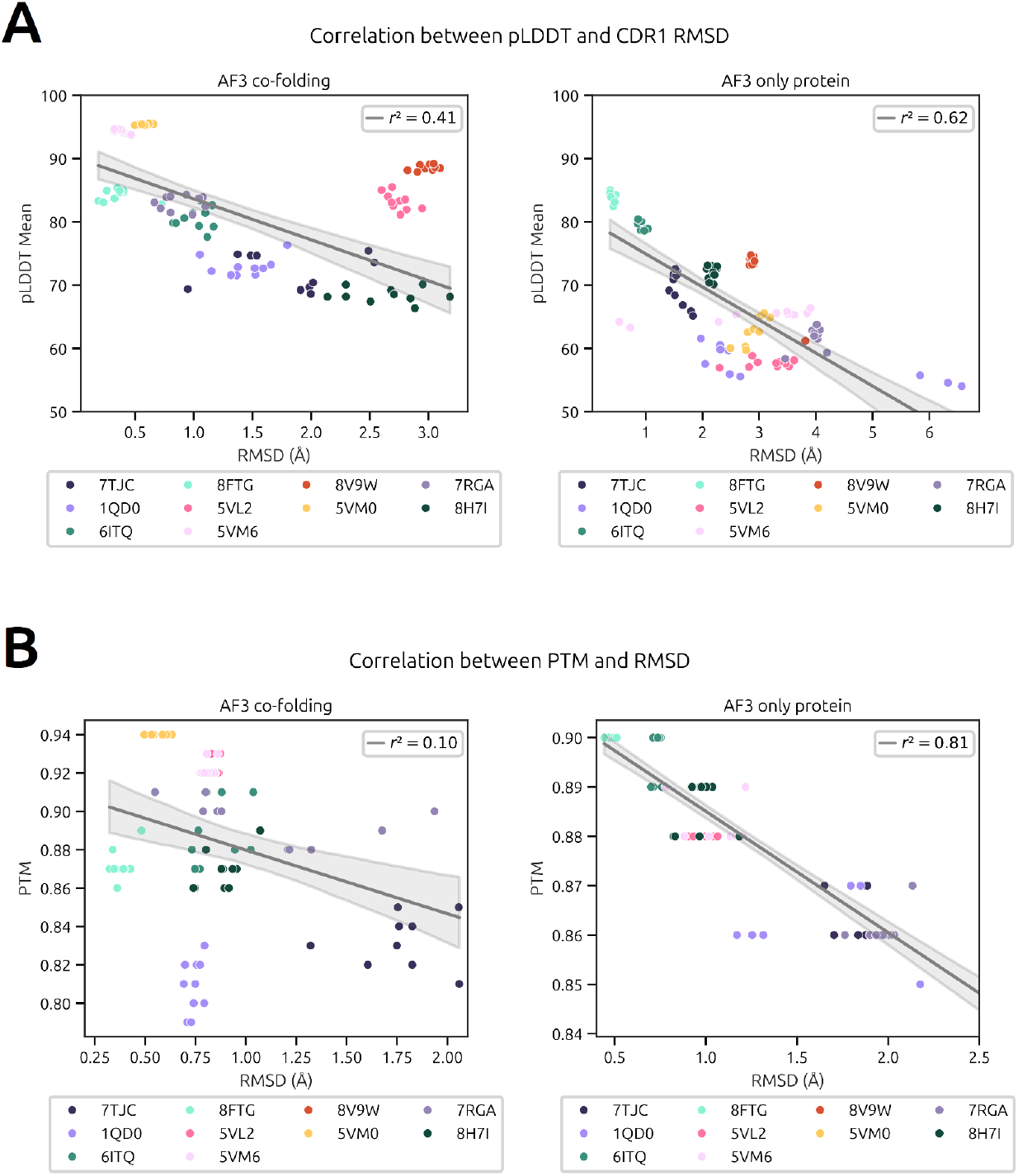
Confidence scores in AF3. **A**: Correlation of CDR pLDDT with CDR RMSD. Points corresponding to PDB 8V9W are considered outliers for the regression. **B**: Correlation of PTM score with nanobody RMSD, for co-folding (left) and monomer (right) predictions.

Furthermore, PTM score correlates with protein RMSD. However, this behaviour was only found for monomer predictions and was lost when including the ligand in the predictions, as can be observed in Figure 5B. This is due to PTM being a measure of confidence of the whole structure. Thus, with monomeric structures it correlates well with Ca RMSD.

Other confidence metrics, like iPTM, have been analysed but no correlation has been found with any structural score (Figure S7), particularly with ligand accuracy measurements.

### Increasing recycles or number of samples does not improve accuracy

Finally, we assess how the accuracy of AF3 and Chai-1 evolves with the number of samples and recycles. These indicate which are the best parameters to obtain the most cost-efficient results. Figures 6 and 7 show the effect of the number of samples and recycles in the predictions of nanobody RMSD, ligand binding site distance and CDR1 RMSD.

**Figure 6:**
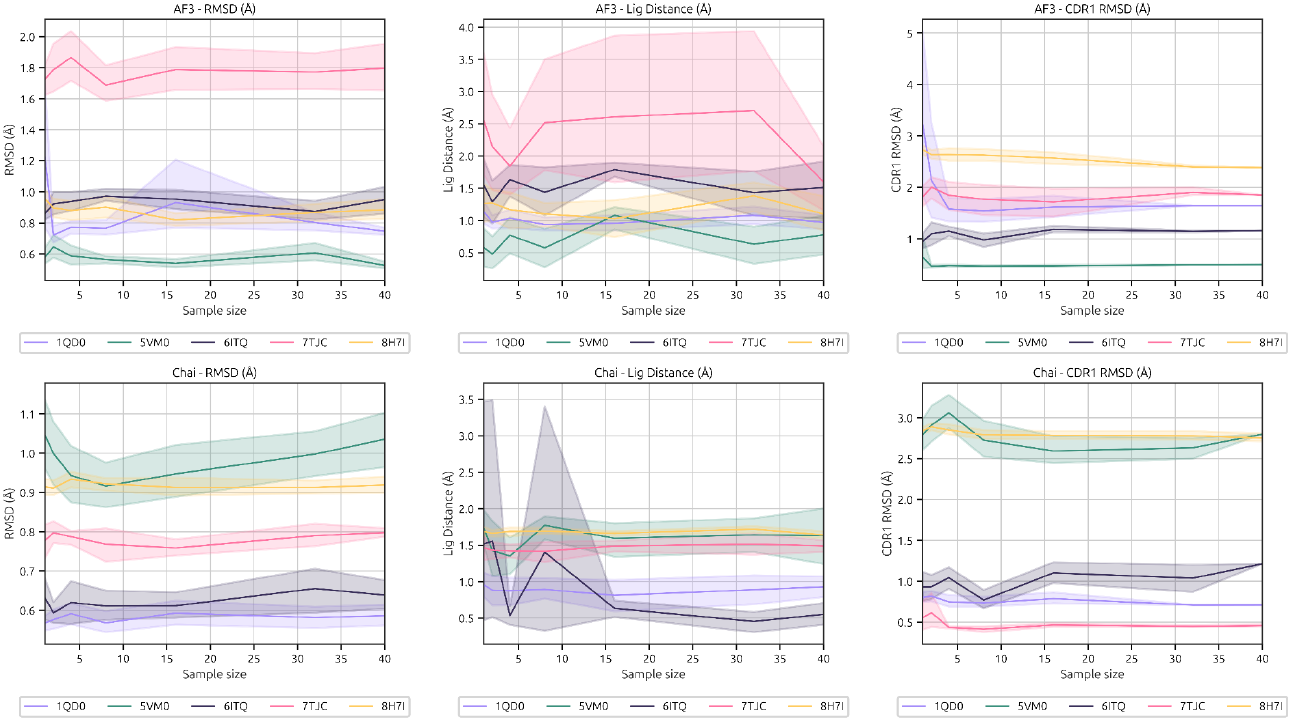
Effect of diffusion sample size on AF3 and Chai-1 accuracy. Results are obtained using the perfect oracle approach, where the best sample for each PDB is considered. From left to right, the plots indicate the effect of sample size on protein, ligand binding site distance, and CDR1 RMSD.

**Figure 7:**
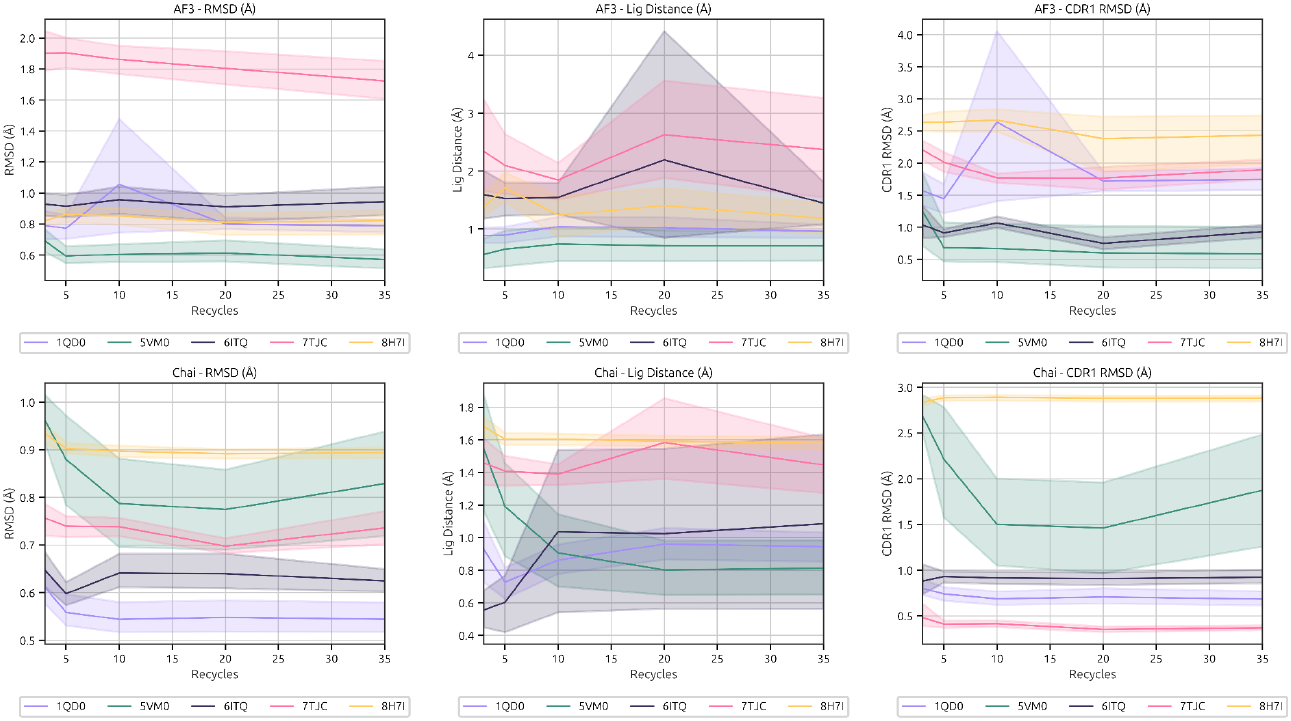
Effect of number of recycles on AF3 and Chai-1 accuracy. Results are obtained using the perfect oracle approach, where the best sample for each PDB is considered. From left to right, the plots indicate the effect of sample size on protein, ligand binding site distance, and CDR1 RMSD.

We wanted to present the results in the most realistic way possible, assessing the most accurate structure only with the provided score of AF3 or Chai-1. However, we found that although the average score of the predicted structures increased with the number of samples, the former did not increase with the number of recycles, indicating that an increase of recycles would not lead to higher agreement with experimental results (Figures S10A and S11A). In addition, we found that for individual PDBs, the aforementioned increase of score does not correlate with an increase in accuracy (Figures S11C and S11C). This is a drawback to blind predictions of sequences, as other metrics would be needed to assess the quality of the obtained structures, convoluting the prediction process.

Following the previous results, we opted to report the “perfect oracle” results, where we show the accuracy for the best possible structure when compared directly with the crystallographic complex. In Figure 6 and Figure S8 we observe that all of the reported metrics stabilize at around the sampling of 10 structures for both Chai-1 and AF3. Therefore, it is a good number of structures to sample, providing a good balance between accuracy and computational cost.

Regarding the number of recycles, previous literature^2,41^ already discussed that a high number of recycles increases the accuracy of the results. However, Figure 7 and Figure S9 show that none of the studied parameters are improved with the number of recycles. If anything, they slightly increase until 5-10 recycles and then stabilizes. This behaviour is seen more profusely in Chai-1 data than in AF3. Hence, increasing the number of recycles above 10 is not recommended, as the computational cost rises without any benefit in structure precision.

## Discussion

Generalized structure predictors, deep learning softwares trained to predict the structure of protein sequences, have shown remarkable accuracy in the modeling of nanobody scaffolds. Higher precision is achieved using co-folding with ligands, probably due to the flexibility of the nanobody ligand-binding region. Among the evaluated predictors, AF3 and Chai-1 outperformed the others in overall structural accuracy, although all models yielded satisfactory results.

However, a significant challenge for structure predictors resides in ligand placement, particularly in ligand orientation. This is a crucial aspect in design pipelines, as accurate ligand placement is essential for identifying key epitope residues for binding. AF3 significantly outperformed other predictors in ligand placement accuracy, highlighting its potential for application in nanobody design workflows. Still, further improvements in the pose of the analyte could be achieved using classic docking softwares.

Further analysis revealed that co-folding predominantly affects the accuracy of CDR1 predictions, whereas CDR2 and CDR3 remain relatively unaffected. Ligand contact analysis indicated a central role for CDR1 in small-molecule binding. This finding is contrasting to protein antigen recognition, typically mediated by CDR3, and suggests a divergence in nanobody binding depending on ligand type. Consequently, nanobody interactions with small molecules and proteins should be studied separately to fully capture these mechanistic differences.

In addition to conventional CDRs, contact analysis uncovered significant contributions to small-molecule binding from the CDR4 region and neighboring framework residues. These findings highlight the importance of considering non-CDR interactions in nanobody design, as binding is not limited to specific regions. Such insights broaden the scope of design strategies by including auxiliary residues that may enhance binding affinity or specificity.

The correlation between pLDDT scores and CDR accuracy reinforces the utility of this metric as an indicator for model confidence in structural predictions. However, the development of a confidence score specifically correlated with ligand placement accuracy would have a more profound impact on design applications.

One of the advantages of current prediction softwares is their relatively low computational cost, enabling accurate predictions with a low number of samples and recycling steps. This efficiency is crucial for design pipelines that require high-throughput sampling of sequence space. However, prediction accuracy was not improved by increasing the number of recycling iterations nor the number of sampled structures.

A notable limitation of this study lies in the small size of the analyzed dataset, constrained by the low availability of nanobody-ligand crystal structures. Despite this, our findings lay important groundwork and highlight the potential of computational structure prediction for nanobody design in the context of small-molecule binding.

## Acknowledgements

The authors thank Albert Ortega-Bartolomé for thoughtful discussions during the development of the paper. ChatGPT (version 5.1) was used to suggest alternative phrasings of some sentences of the manuscript, but never adding to or changing the meaning of the original text by the authors.

## Methods

### Nanobody - analyte dataset

The Nb-analyte dataset has been curated using the following approach: First, SAbDab^32^ was queried using the keywords “Nanobody + hapten”, prompting the search for all the structures containing nanobody strands targeting a small molecule analyte. From there, 69 protein chains were obtained, with their CDRs. They were then filtered using pandas^42^ to procure only the chains that contained unique CDRs, as the complementary regions have higher amino acid variability and are tailored for binding. Mutations in framework regions were not considered significant for the binding process and thus would probably not lead to changes in structure worth analysing.

After that filtering, 12 Nb-SM systems were left and upon further visual inspection with PyMOL the structure corresponding to 7TE8^43^ was discarded. It corresponded to a dual biosensor, where two nanobodies with different paratopes bound to a small molecule. This was not the geometry we were searching for and differs enough from the other nanobodies in the dataset to be analysed and processed differently.

From the final dataset, 5 structures were published in the PDB prior to 2021, and thus would be included in most of the training sets for the structure predictors analysed in this paper. Those complexes are 1QD0^44^, 5VL2, 5VM0 and 5VM6^36^; and 6ITQ^45^. The other remaining structures that were not included in the training sets were 7TJC^46^, 8FTG^47^, 8V9W^39^, 7RGA^48^, and 8H7I^38^ (Table S1).

### Input generation and structure pre-processing

To generate the inputs for the structure predictions, an automated pipeline was used, available at https://github.com/QTC-IQAC/predictorVerse. It allowed us to generate the necessary inputs with just the protein sequence and ligand SMILES. The protein sequences were obtained from SAbDab, and the ligand SMILES were obtained by loading the PDB structure with MDTraj^49^, separating the protein and the ligand chains and extracting the SMILES code from those small molecules using RDKit^50^. However, some automatically generated SMILES were confusing for some structure predictors and gave erroneous results, be it not interpreting the molecule at all or providing the wrong stereochemistry. Thus, we manually replaced those SMILES for the ones defined in PubChem for the associated analytes. This has been done for the structures 7RGA and 8H7I. For 1QD0, the copper atoms have been left out of the predictions.

### Structure predictions

The results have been obtained using the aforementioned structure predictors: AF3^2^, Chai-1^3^, Boltz-1 and Boltz-1x^4^, RFAA^33^, FlowDock^35^ and OmegaFold^34^. We decided to include FlowDock, with and without using a template from Chai-1, as its affinity predictions could provide reliable insight in a protein design pipeline. OmegaFold was included as a previous benchmark reported it yielded good results for the modelling of antibodies and nanobodies^51^. However, it is not capable of producing co-folding predictions (see Table 1).

**Table 1:** Information about the protein folding capabilities of the analysed structure predictors.

|  | AF3 | Chai-1 | Boltz-1/1x | RFAA | FlowDock | OmegaFold |
| --- | --- | --- | --- | --- | --- | --- |
| Predicts only protein? | YES | YES | YES | YES | NO | YES |
| Predicts protein + ligand (co-folding)? | YES | YES | YES | YES | YES | NO |

For general results they have been run with the default hyper-parameters and 10 samples obtained. For the case of RFAA and OmegaFold, only 1 sample per nanobody structure was considered as there were no specific options to sample more structures. Predictions were run on several computational resources, using Nvidia GeForce RTX™ 4070 SUPER, Nvidia GeForce RTX™ 4090 and Nvidia H100 GPUs.

### Analysis

Prior to the analysis, the original PDBs were cleaned out to be easier to manipulate. Any extra chains were removed to only leave one instance of nanobody with its corresponding small molecule. The water atoms were also removed.

Analysis of the structures was performed using MDTraj^49^, pandas^52^, seaborn^53,54^ and NumPy^55^. For the analysis of contacts, we added hydrogens to the ligand with PyMOL^56^ and to the protein with PDBFixer^57,58^. Then the contacts were assessed using PLIP^59^. We only considered the neighbourhoods for CDR1 and CDR3 as they are the largest CDRs and their surrounding residues might overlap with the neighbour regions of both CDR2 and the CDR4 region.

