## Supplementary figures and tables for "Assessing Structural Prediction Accuracy for Nanobody–Small Molecule Complexes"

### Supporting Information

**Table S1**: Information about the nanobody-small molecule complexes found in SAb-Dab. Complexes published in PDB before 2021 are in the training sets of AF3, Chai-1, Boltz-1. Boltz-1x and RFAA.

| PDB ID | Publication Date | In PDBBind? | In UniRef? | Comments | Analyte Name | Analyte Structure |
|---|---|---|---|---|---|---|
| **1QD0** | 2000 | NO | NO | | RR6 | 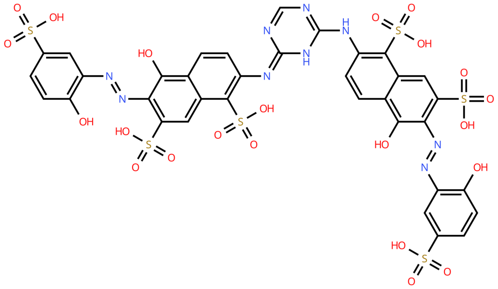 |
| **5VL2** | 2018 | YES | NO | | triclocarban | 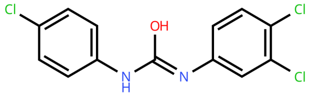 |
| **5VM6** | 2018 | YES | NO | | triclocarban | 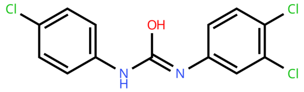 |
| **5VM0** | 2018 | YES | NO | | triclocarban | 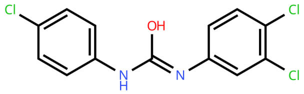 |
| **6ITQ** | 2019 | YES | NO | | cortisol | 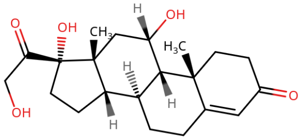 |
| **7TJC** | 2022 | NO | NO | | chloramphenicol | 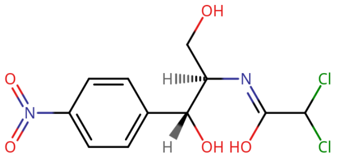 |

| | | | | | | |
|---|---|---|---|---|---|---|
| **7RGA** | 2022 | NO | NO | | methotrexate | 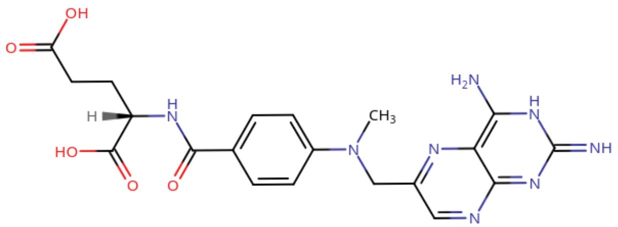 |
| **8FTG** | 2023 | NO | NO | | caffeine | 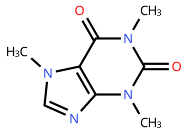 |
| **8H7I** | 2023 | NO | NO | | quinalphos | 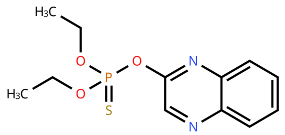 |
| **8V9W** | 2024 | NO | NO | | fentanyl | 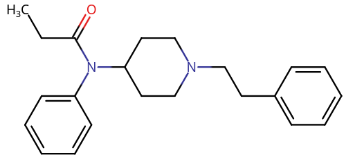 |
| **7TE8** | 2022 | NO | NO | Dual Biosensor. Excluded | cannabidiol | 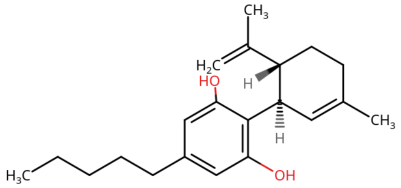 |

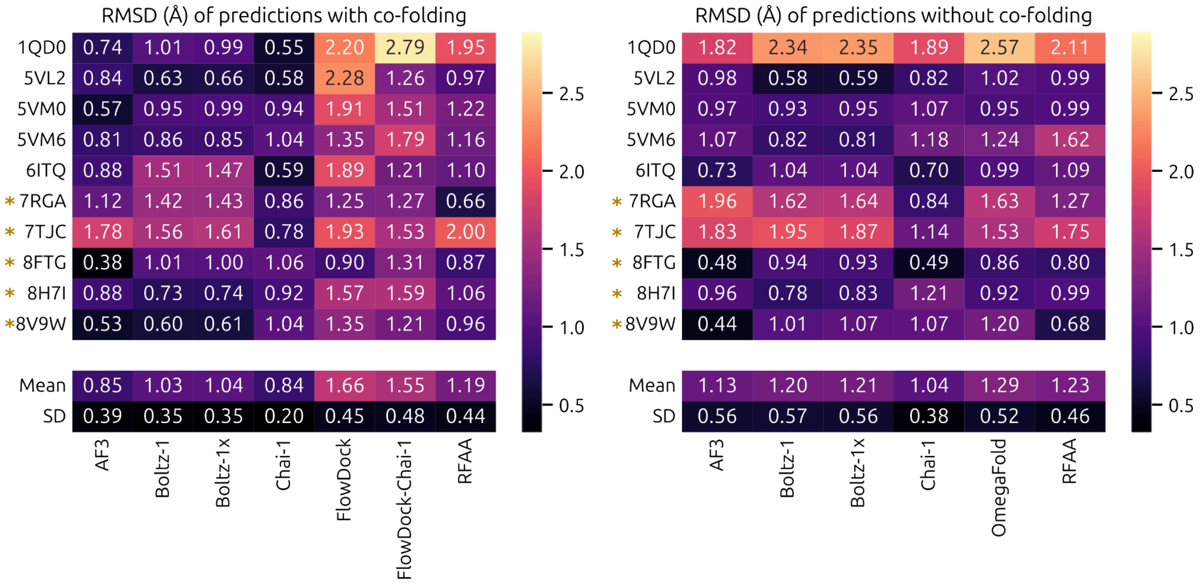


**Figure S1: Overview of nanobody RMSD for each complex and predictor.** * indicates that the system is not in the training set of the predictors. **A**: Results obtained for co-folding predictions **B**: Results for monomer predictions.

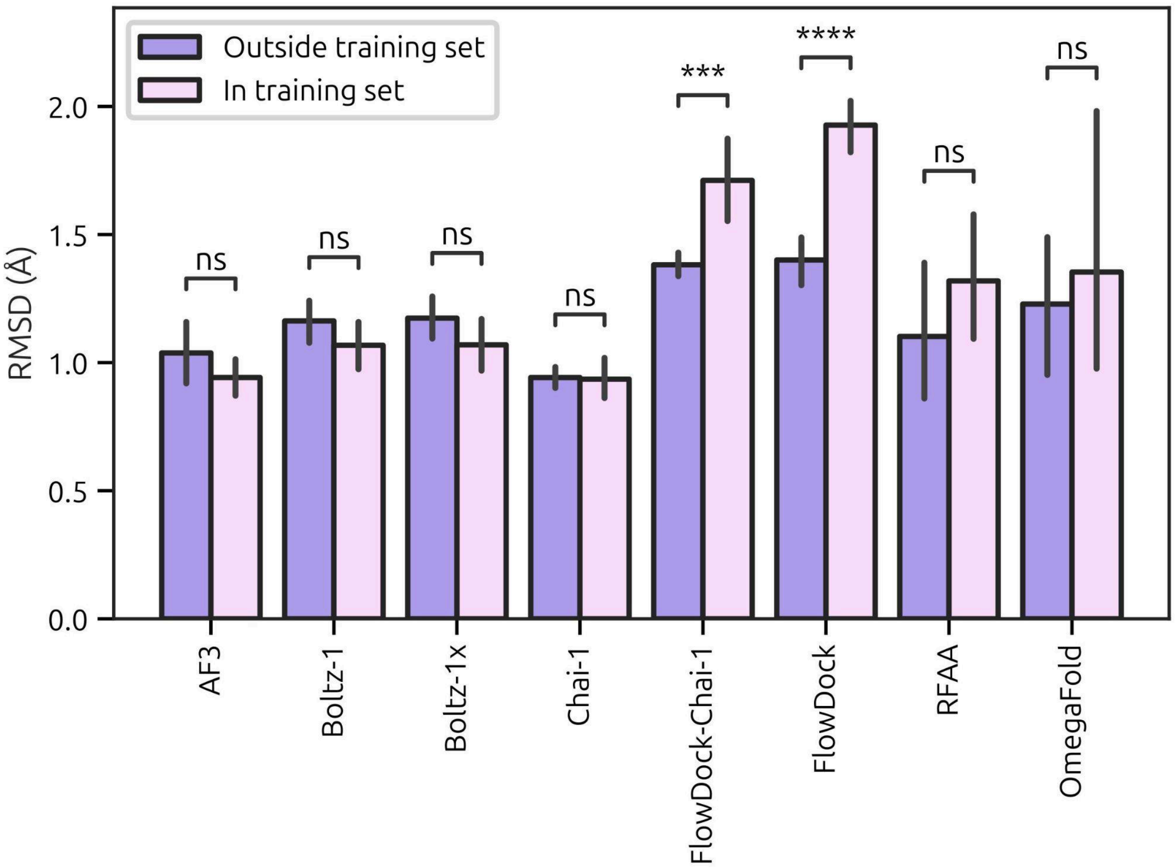


**Figure S2: Effect of training set in the predictions of nanobody structure, accounting for both co-folding and only protein structures.**

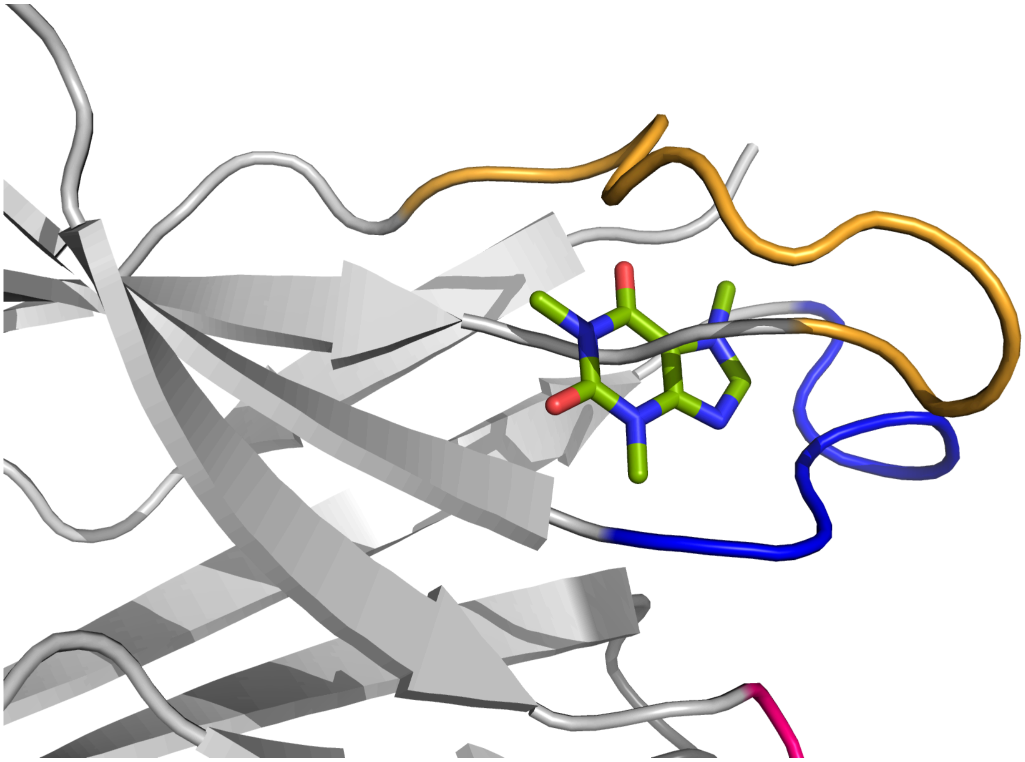

**Figure S3: RFAA binding.** An example of the steric clashes that RFAA produced when generating complex predictions. The displayed nanobody is 8FTG, binding caffeine.

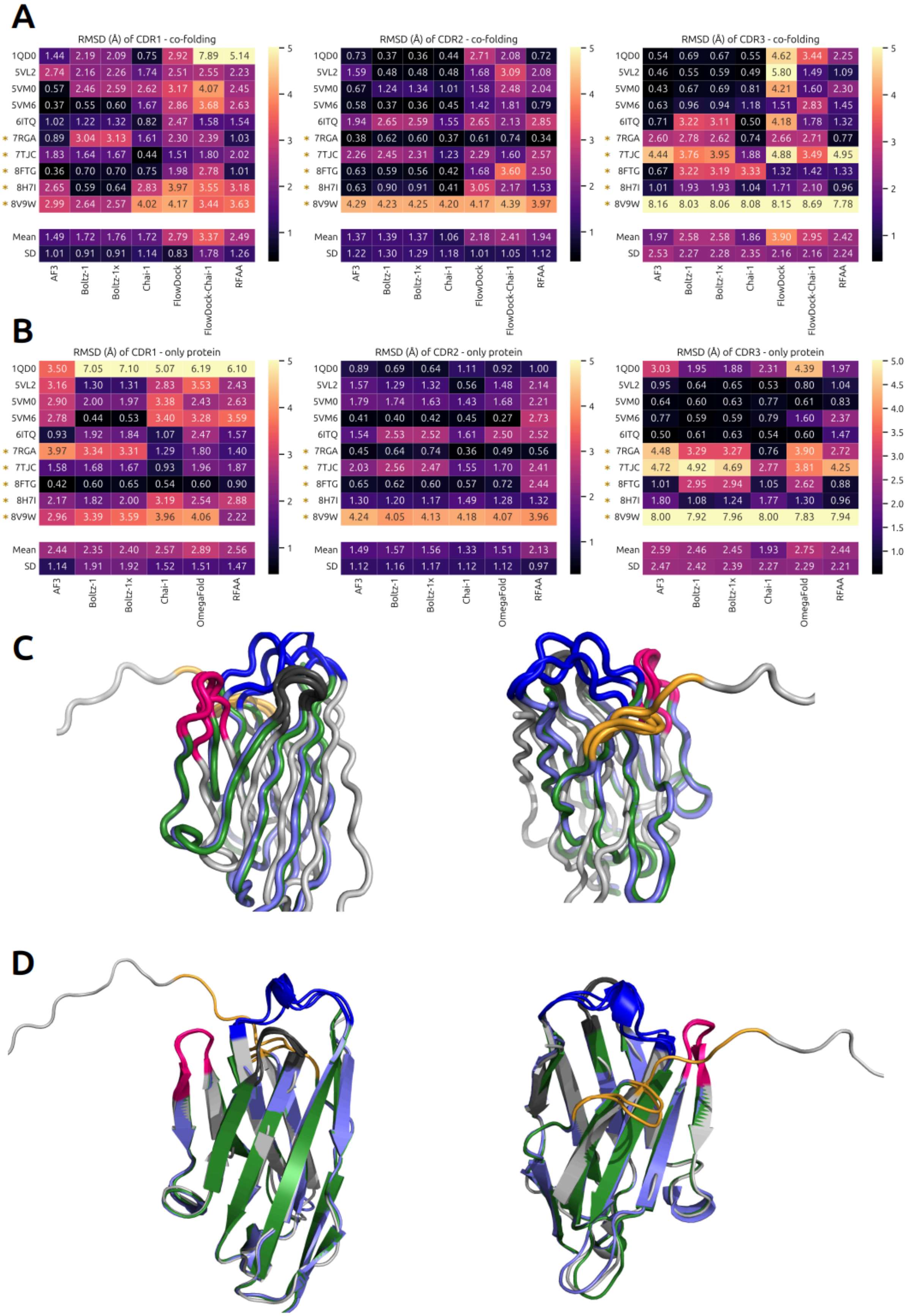
A
RMSD (Å) of CDR1 - co-folding
1QD0 1.44 2.19 2.09 0.75 2.92 7.89 5.14
5VL2 2.74 2.16 2.26 1.74 2.51 2.55 2.23
5VM0 0.57 2.46 2.59 2.62 3.17 4.07 2.45
5VM6 0.37 0.55 0.60 1.67 2.86 3.68 2.63
6ITQ 1.02 1.22 1.32 0.82 2.47 1.58 1.54
* 7RGA 0.89 3.04 3.13 1.61 2.30 2.39 1.03
* 7TJC 1.83 1.64 1.67 0.44 1.51 1.80 2.02
* 8FTG 0.36 0.70 0.70 0.75 1.98 2.78 1.01
* 8H7I 2.65 0.59 0.64 2.83 3.97 3.55 3.18
* 8V9W 2.99 2.64 2.57 4.02 4.17 3.44 3.63
Mean 1.49 1.72 1.76 1.72 2.79 3.37 2.49
SD 1.01 0.91 0.91 1.14 0.83 1.78 1.26
AF3 Boltz-1 Boltz-1x Chai-1 FlowDock FlowDock-Chai-1 RFAA
RMSD (Å) of CDR2 - co-folding
1QD0 0.73 0.37 0.36 0.44 2.71 2.08 0.72
5VL2 1.59 0.48 0.48 0.48 1.68 3.09 2.08
5VM0 0.67 1.24 1.34 1.01 1.58 2.48 2.04
5VM6 0.58 0.37 0.36 0.45 1.42 1.81 0.79
6ITQ 1.94 2.65 2.59 1.55 2.65 2.13 2.85
* 7RGA 0.38 0.62 0.60 0.37 0.61 0.74 0.34
* 7TJC 2.26 2.45 2.31 1.23 2.29 1.60 2.57
* 8FTG 0.63 0.59 0.56 0.42 1.68 3.60 2.50
* 8H7I 0.63 0.90 0.91 0.41 3.05 2.17 1.53
* 8V9W 4.29 4.23 4.25 4.20 4.17 4.39 3.97
Mean 1.37 1.39 1.37 1.06 2.18 2.41 1.94
SD 1.22 1.30 1.29 1.18 1.01 1.05 1.12
AF3 Boltz-1 Boltz-1x Chai-1 FlowDock FlowDock-Chai-1 RFAA
RMSD (Å) of CDR3 - co-folding
1QD0 0.54 0.69 0.67 0.55 4.62 3.44 2.25
5VL2 0.46 0.55 0.59 0.49 5.80 1.49 1.09
5VM0 0.43 0.67 0.69 0.81 4.21 1.60 2.30
5VM6 0.63 0.96 0.94 1.18 1.51 2.83 1.45
6ITQ 0.71 3.22 3.11 0.50 4.18 1.78 1.32
* 7RGA 2.60 2.78 2.62 0.74 2.66 2.71 0.77
* 7TJC 4.44 3.76 3.95 1.88 4.88 3.49 4.95
* 8FTG 0.67 3.22 3.19 3.33 1.32 1.42 1.33
* 8H7I 1.01 1.93 1.93 1.04 1.71 2.10 0.96
* 8V9W 8.16 8.03 8.06 8.08 8.15 8.69 7.78
Mean 1.97 2.58 2.58 1.86 3.90 2.95 2.42
SD 2.53 2.27 2.28 2.35 2.16 2.16 2.24
AF3 Boltz-1 Boltz-1x Chai-1 FlowDock FlowDock-Chai-1 RFAA
1 2 3 4 5
B
RMSD (Å) of CDR1 - only protein
1QD0 3.50 7.05 7.10 5.07 6.19 6.10
5VL2 3.16 1.30 1.31 2.83 3.53 2.43
5VM0 2.90 2.00 1.97 3.38 2.43 2.63
5VM6 2.78 0.44 0.53 3.40 3.28 3.59
6ITQ 0.93 1.92 1.84 1.07 2.47 1.57
* 7RGA 3.97 3.34 3.31 1.29 1.80 1.40
* 7TJC 1.58 1.68 1.67 0.93 1.96 1.87
* 8FTG 0.42 0.60 0.65 0.54 0.60 0.90
* 8H7I 2.17 1.82 2.00 3.19 2.54 2.88
* 8V9W 2.96 3.39 3.59 3.96 4.06 2.22
Mean 2.44 2.35 2.40 2.57 2.89 2.56
SD 1.14 1.91 1.92 1.52 1.51 1.47
AF3 Boltz-1 Boltz-1x Chai-1 OmegaFold RFAA
RMSD (Å) of CDR2 - only protein
1QD0 0.89 0.69 0.64 1.11 0.92 1.00
5VL2 1.57 1.29 1.32 0.56 1.48 2.14
5VM0 1.79 1.74 1.63 1.43 1.68 2.21
5VM6 0.41 0.40 0.42 0.45 0.27 2.73
6ITQ 1.54 2.53 2.52 1.61 2.50 2.52
* 7RGA 0.45 0.64 0.74 0.36 0.49 0.56
* 7TJC 2.03 2.56 2.47 1.55 1.70 2.41
* 8FTG 0.65 0.62 0.60 0.57 0.72 2.44
* 8H7I 1.30 1.20 1.17 1.49 1.28 1.32
* 8V9W 4.24 4.05 4.13 4.18 4.07 3.96
Mean 1.49 1.57 1.56 1.33 1.51 2.13
SD 1.12 1.16 1.17 1.12 1.12 0.97
AF3 Boltz-1 Boltz-1x Chai-1 OmegaFold RFAA
RMSD (Å) of CDR3 - only protein
1QD0 3.03 1.95 1.88 2.31 4.39 1.97
5VL2 0.95 0.64 0.65 0.53 0.80 1.04
5VM0 0.64 0.60 0.63 0.77 0.61 0.83
5VM6 0.77 0.59 0.59 0.79 1.60 2.37
6ITQ 0.50 0.61 0.63 0.54 0.60 1.47
* 7RGA 4.48 3.29 3.27 0.76 3.90 2.72
* 7TJC 4.72 4.92 4.69 2.77 3.81 4.25
* 8FTG 1.01 2.95 2.94 1.05 2.62 0.88
* 8H7I 1.80 1.08 1.24 1.77 1.30 0.96
* 8V9W 8.00 7.92 7.96 8.00 7.83 7.94
Mean 2.59 2.46 2.45 1.93 2.75 2.44
SD 2.47 2.42 2.39 2.27 2.29 2.21
AF3 Boltz-1 Boltz-1x Chai-1 OmegaFold RFAA
1.0 1.5 2.0 2.5 3.0 3.5 4.0 4.5 5.0
C
D

**Figure S4: CDR RMSD for each nanobody and predictor.** * indicates that the system is not in the training set of the predictors. **A**: Heatmaps for co-folding predictions. **B**: Heatmap for monomer predictions. For both A and B, the RMSD of CDR2 and CDR3 for 8V9W is abnormally high compared to compounds. This is due to a misalignment from MDTraj. This is due to the long C-terminal coil that this nanobody has. MDTRaj tries to minimize the overall RMSD, greatly affected by the outlier tail at the cost of the alignment of other residues. Other alignment tools, like the one in PyMOL prioritize the alignment of the core nanobody instead of the C-terminal coil. The other cases with high CDR RMSD values correspond to differences in the predicted and crystallographic structures, as the alignments of MDTraj and PyMOL are equal. **C**: MDTraj alignment for 8V9W, Cα only. **D**: PyMOL alignment for 8V9W. For both C and D: Crystal structure in white, AF3 in green and Chai in purple. CDR1 in blue, CDR2 in pink and CDR3 in orange.

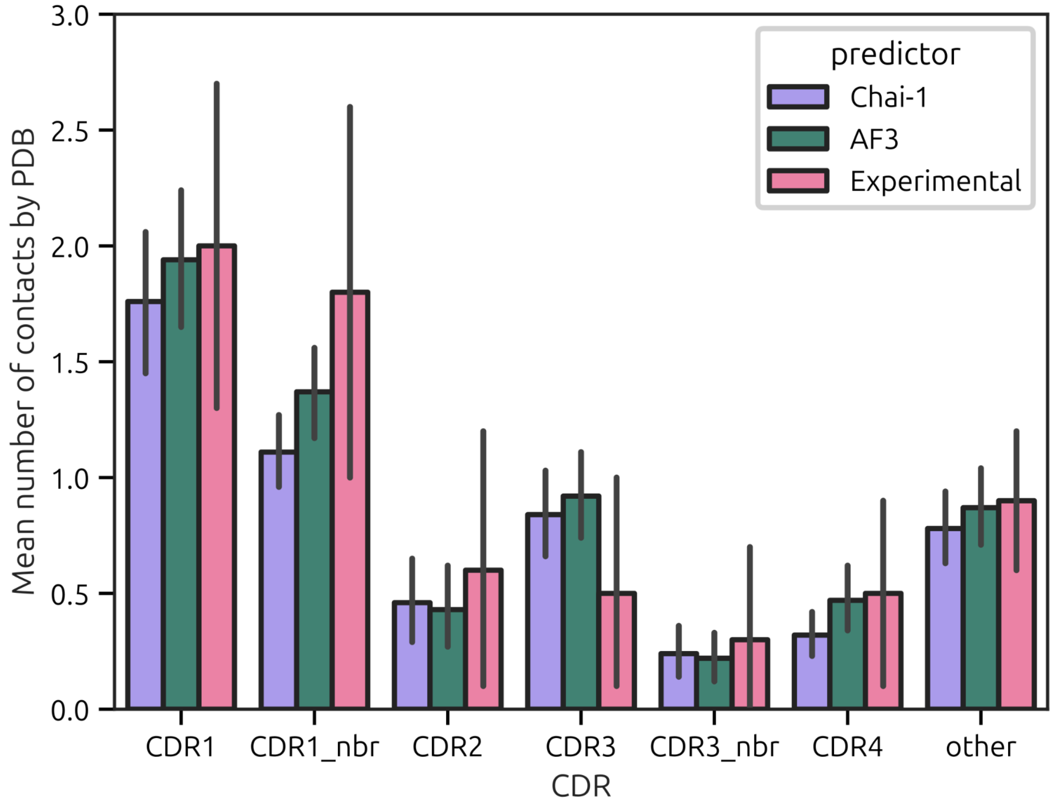


**Figure S5: Comparison of the number of contacts between AF3, Chai-1 and crystallographic structures, for CDRs and its neighbourhood residues.**

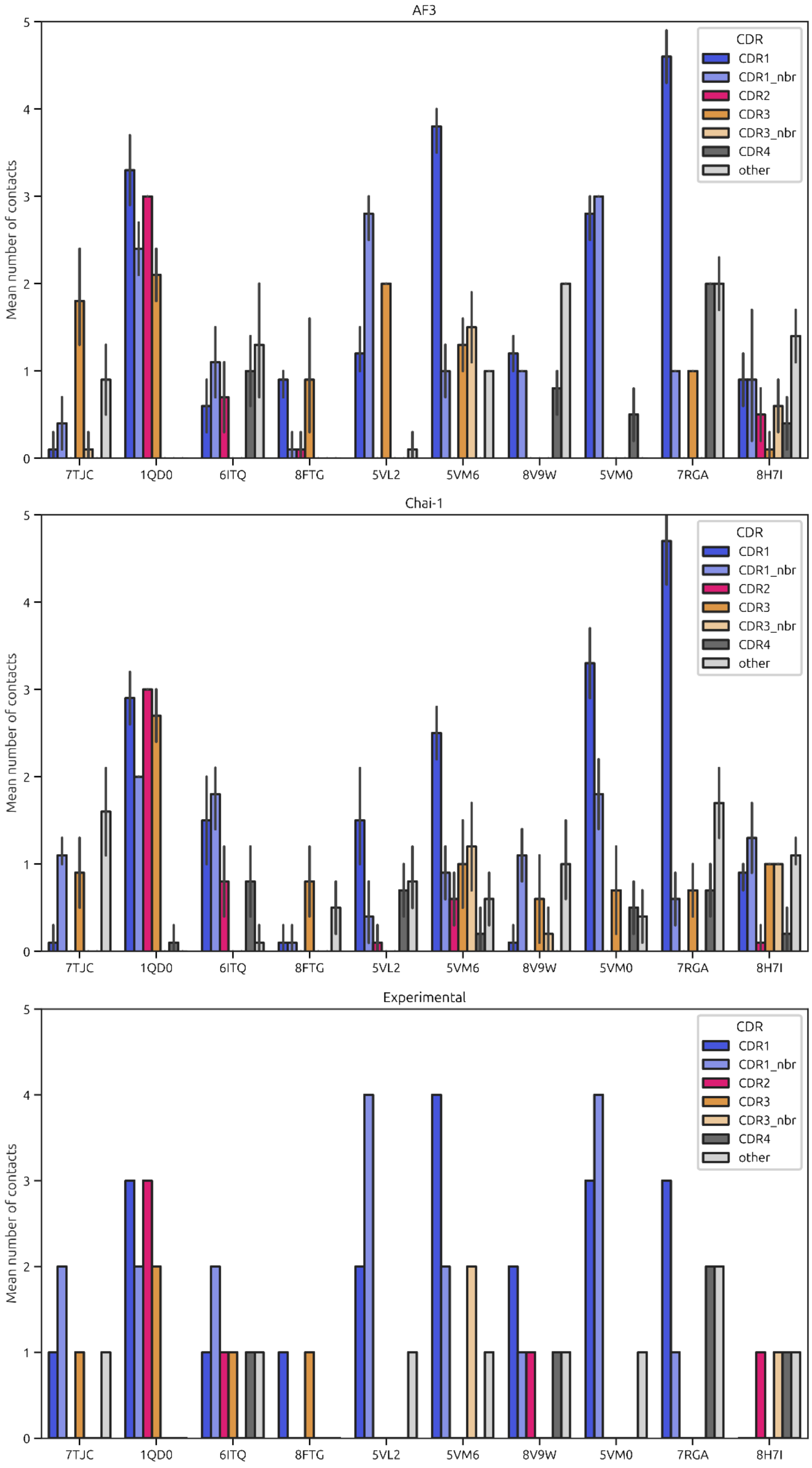
AF3
Chai-1
Experimental
Mean number of contacts
0
1
2
3
4
5
CDR
CDR1
CDR1_nbr
CDR2
CDR3
CDR3_nbr
CDR4
other
7TJC
1QD0
6ITQ
8FTG
5VL2
5VM6
8V9W
5VM0
7RGA
8H7I

**Figure S6: Number of CDR and neighbourhood residues contacts by complex. Results for AF3, Chai-1 and crystallographic structures.**

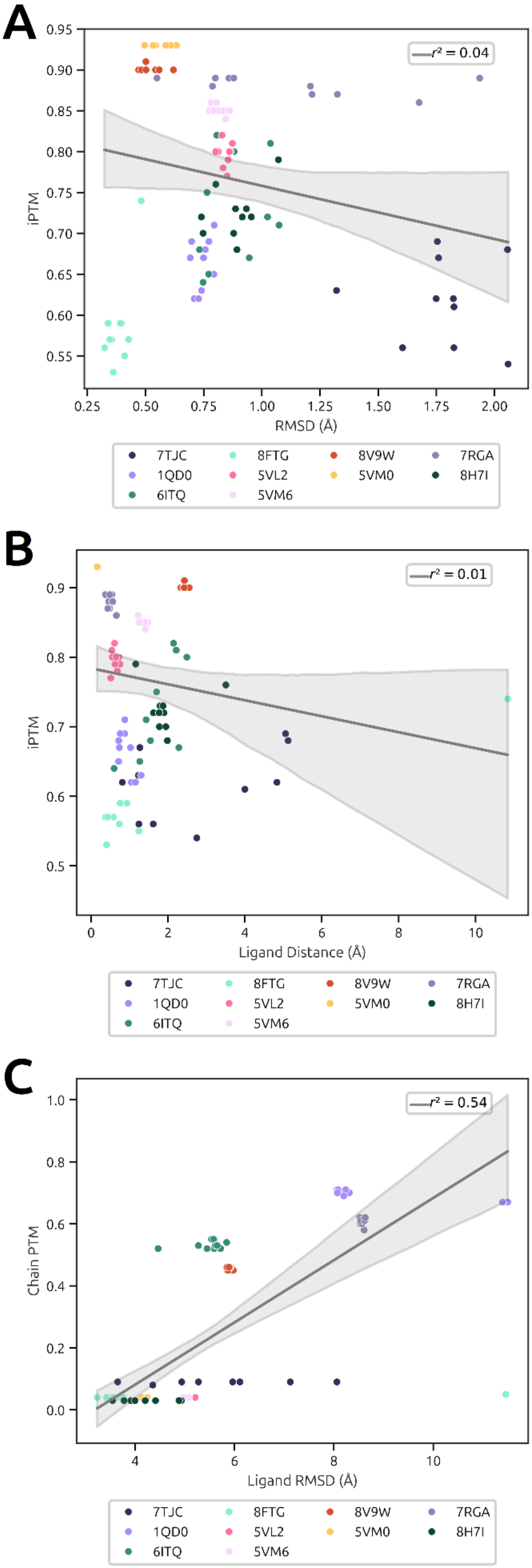
A
B
C
r² = 0.04
r² = 0.01
r² = 0.54
iPTM
RMSD (Å)
Ligand Distance (Å)
Chain PTM
Ligand RMSD (Å)
7TJC
8FTG
8V9W
7RGA
1QD0
5VL2
5VM0
8H7I
6ITQ
5VM6

**Figure S7: Correlation between AF3 scores and ligand accuracy measures.** There is no correlation between iPTM score and **A:** ligand RMSD**, B:** ligand binding site distance**. C:** Chain PTM score for the ligand chain partially correlates with ligand RMSD**.**

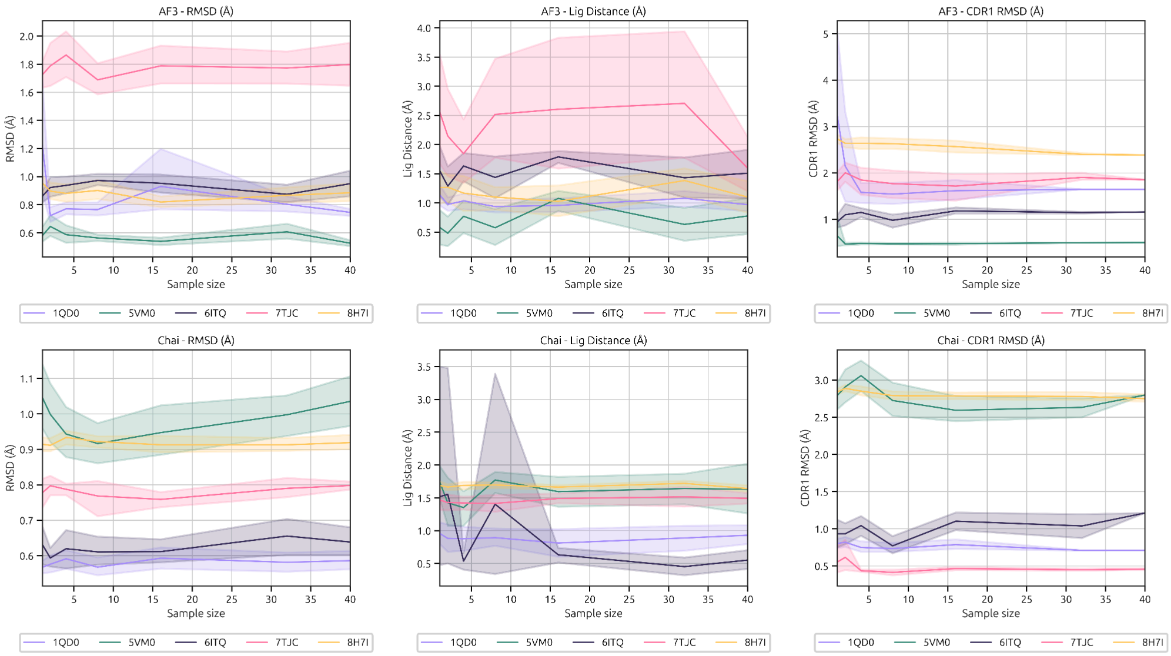


**Figure S8**: **Effect of diffusion sample size on AF3 and Chai-1 accuracy.** Results are obtained using the perfect oracle approach, where the best sample for each PDB is considered. From left to right, the plots indicate the effect of sample size on protein, ligand binding site distance, and CDR1 RMSD.

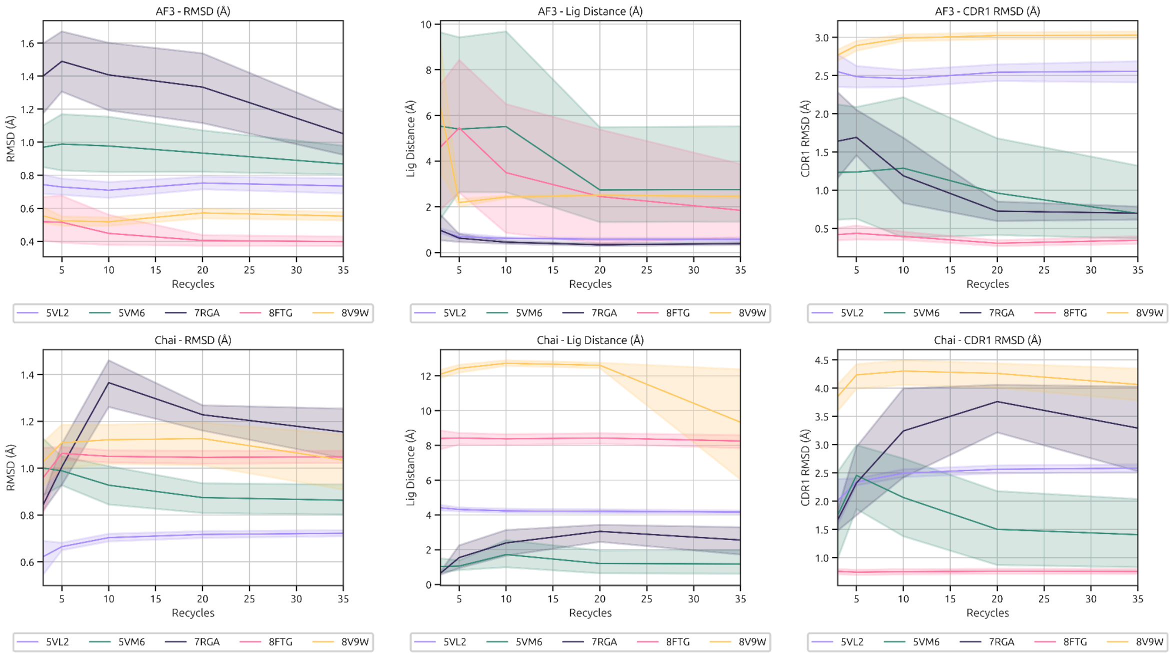


**Figure S9**: **Effect of number of recycles on AF3 and Chai-1 accuracy.** Results are obtained using the perfect oracle approach, where the best sample for each PDB is considered. From left to right, the plots indicate the effect of sample size on protein, ligand binding site distance, and CDR1 RMSD.

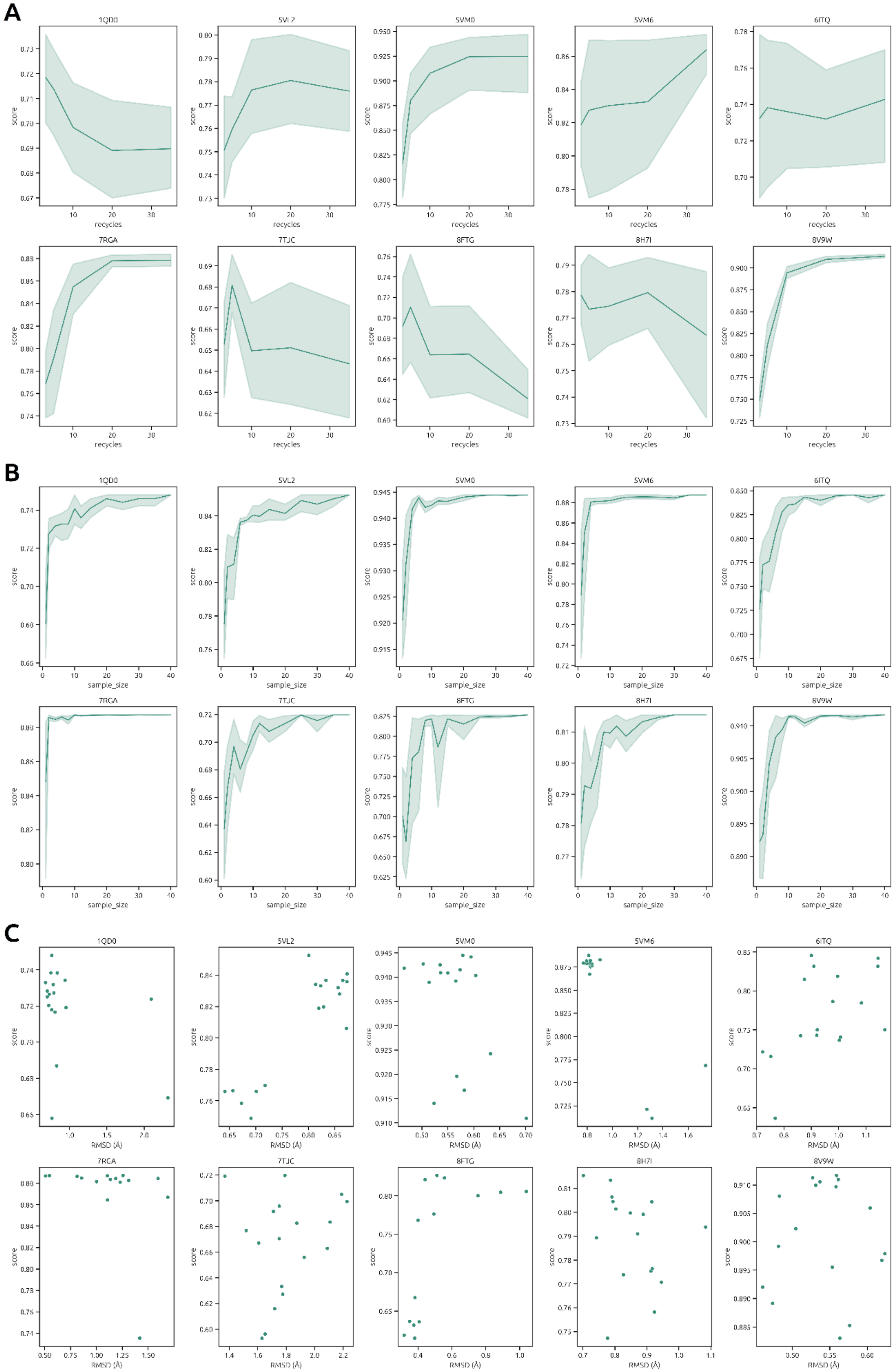
A
1QD0
5VL2
5VM0
5VM6
6ITQ
7RGA
7TJC
8FTG
8H7I
8V9W
score
recycles
B
sample_size
C
RMSD (Å)

**Figure S10**: **Correlation of AF3 score with sample size and recycles, for each complex. A**: There is no improvement in score with an increase of recycles, for each individual complex. **B:** The score increases with the number of samples, reaching a maximum at around 10 samples. **C**: Score does not correlate with RMSD for individual complexes.

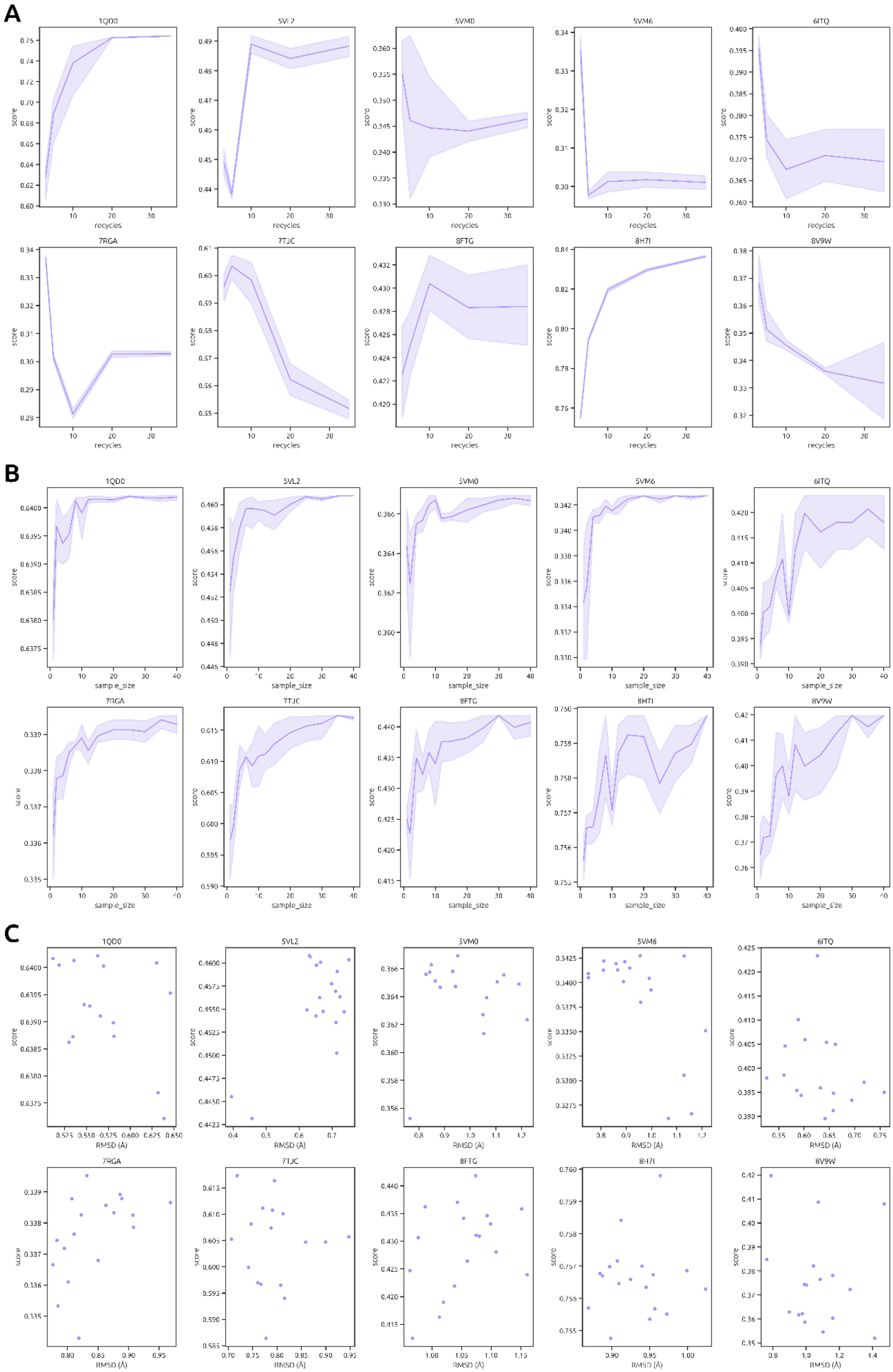
A
1QD0
5VL2
5VM0
5VM6
6ITQ
7RGA
7TJC
8FTG
8H7I
8V9W
score
recycles
B
sample_size
C
RMSD (Å)

**Figure S11**: **Correlation of Chai-1 score with sample size and recycles, for each complex.** A: There is no improvement in score with an increase of recycles, for each individual complex. **B:** The score increases with the number of samples, reaching a maximum at around 10 samples. **C**: Score does not correlate with RMSD for individual complexes.
